# A biologically realistic model to predict wildlife-vehicle collision risks

**DOI:** 10.1101/2023.02.15.528614

**Authors:** Annaëlle Bénard, Thierry Lengagne, Christophe Bonenfant

## Abstract

Road networks have major ecological impacts on living organisms consequent to habitat loss and fragmentation, chemical and acoustic pollution, and direct mortality when wildlife-vehicle collisions are involved (WVC). The many past empirical studies revealed biological traits shared by species most vulnerable to roadkills (e.g. population density). Similarly, spatial locations of WVC hot-spots are associated to landscape features at large spatial scales, and to road characteristics at small spatial scale. We currently lack a comprehensive theoretical framework for WVC. Animal movement in relation to habitats is an essential driver of encounters with roads, but this remains largely ignored in studies, even when movement ecology provides the necessary tools to investigate the impact of animal movement on WVC. We built a general individual-based model incorporating recent knowledge in movement ecology (movement typology: roaming, migratory route crossing a road, active attraction and active repulsion of roads) to estimate WVC risks. We explored the relative effects of animal and vehicle movement parameters (speed, abundance, road sinuosity and animal movement pattern) on collision probability. We show that animal behaviour toward roads has major impacts on the number and risks of WVC, but also modulate the effects of other factors (animal traveling speed, species local abundance, road traffic volume) on WVC. Sensitivity analyses show that the movement and behaviour of the animal has more influence on WVC risks than any of the characteristics of roads and vehicles we tested. Our results suggest that (1) effective roadkill mitigation should be species-specific and could vary in efficiency depending on the target’s movement pattern (mating and migratory seasons, foraging habits…) and (2) empirical studies of WVC should incorporate knowledge about the behavioural habits of the focal species in relation to roads.

## 1 Introduction

The last century has witnessed a continuous and tremendous urbanisation of almost all ecosystems with profound influences on living organisms (Corlett, 2015). A particularly important anthropogenic factor generating major ecosystem modification and disturbance is the increasing densification and encroachment of the earth-bound transportation systems, and particularly of the road network (Davenport & Davenport, 2006). The presence of roads modify the local and large-scale environment of species as a consequence of chemical and noise pollution, changes in vegetation diversity, habitat loss and fragmentation (see Forman & Alexander, 1998; Coffin, 2007, for reviews). Roads can also have profound effects on ecosystem functioning and the biology of living organisms (Holderegger & Di Giulio, 2010). For instance, road mortality is a key driver of biodiversity loss in amphibians (Hels & Buchwald, 2001), birds and mammals (Grilo *et al*., 2020) and has been shown to affect animal behaviours (Andrews & Gibbons, 2005; Legagneux & Ducatez, 2013). In the context of the on-going biodiversity crisis (Ceballos & Ehrlich, 2002), knowledge about the mechanisms that lead to road mortality in wildlife is therefore critical.

For decades, empirical studies have been accumulating, reporting on collision hot-spots (Teixeira *et al*., 2013; Silveira Miranda *et al*., 2020), animal foraging habits near roadsides (Carvalho-Roel *et al*., 2019), or the seasonal variation in the number of reported cases (Medinas *et al*., 2021). We now have a clearer picture of what factors mainly account for collision risks (Table 1). Species with life history traits such as extended home ranges or slow travel speeds, and species occurring at high abundances are more often subjected to roadkill. Road characteristics (such as high traffic volume) and landscape features (*e.g.* habitats promoting high levels of animal activity and mobility) account for WVC hot-spots locations at small spatial scales, but collisions distribution can also be predicted for large spatial scales with variables such as road density, hunting bags (Saint-Andrieux *et al*., 2020) or landscape connectivity (Girardet *et al*., 2015). However, the relative effects of risk factors are difficult to disentangle. For example, as traffic mortality depletes local populations, roadkill numbers will decrease and simply counting collisions in high-risk areas could lead to an underestimation of the importance of animal abundance as a key determining factor of roadkill hot-spots (Teixeira *et al*., 2017). So far few attempts at a theoretical model for wildlife-vehicle collisions (WVC) have been made, and our current knowledge on the quantitative effects of traffic on encounter and mortality risks for wildlife remains essentially descriptive, with little predictive abilities.

**Table 1:**
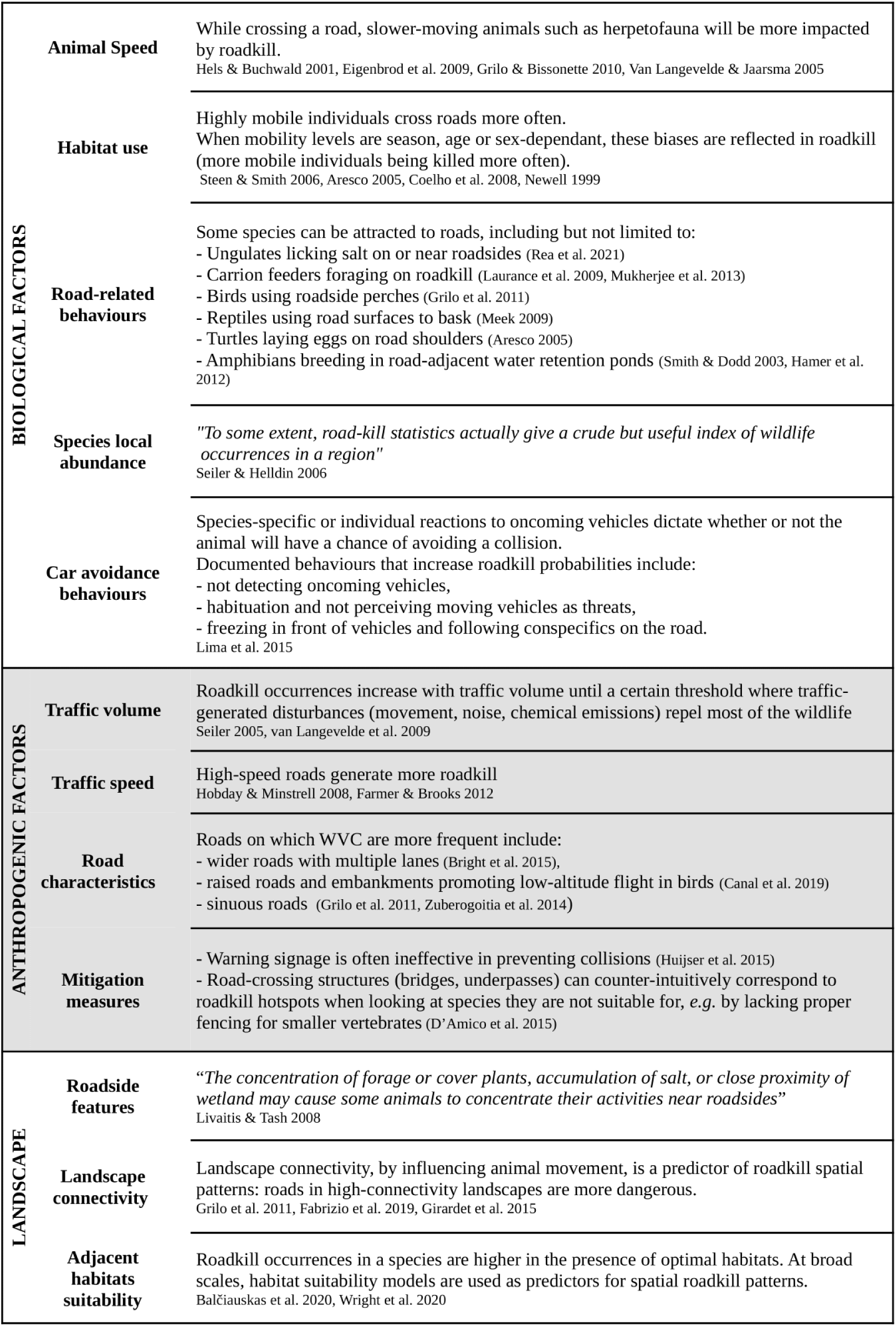
An overview of the different factors correlated to the incidence of wildlife-vehicle collisions in the current literature.

The very first model dealing with collision probabilities at a large spatial scale is the ideal gas model, first coined by physicist J. C. Maxwell in the context of molecule collisions in a gas (Maxwell, 1860). The ideal gas model posits that encounter rate within a defined volume or plane is a function of molecule speed, density and collision zone (Hutchinson & Waser, 2007). Despite its heuristic value in ecology, to estimate encounter rates between males and females of a given species for example, it has seldom been used to study WVC as the general application in this context is not straightforward. For example, in most wildlife-vehicle encounters the animal is likely to die and will no longer be at risk of collision again, which is not reflected in colliding molecules. Others proposed WVC-specific mechanistic models: Hels & Buchwald (2001) modelled the probability of death for an amphibian crossing a road (see Gibbs & Shriver, 2002; van Langevelde & Jaarsma, 2005, for an application to other species), allowing the integration of road mortality into population persistence models (Gibbs & Shriver, 2002).

A shortcoming shared by all current WVC models is the lack of consideration for animal movement and habitat use behaviour, often limiting when one aims at describing the complexity of animal movement in an heterogeneous landscape (Siniff & Jessen, 1969). For example, not only do animals cross roads, but they can roam along roadsides to forage hence putting them at much higher risk of being hit by a car (scavenging: Schwartz *et al*. 2018; Ratton *et al*. 2014, hunting: Gomes *et al*. 2009; D’Amico *et al*. 2013, salt licking: Fraser & Thomas 1982). More importantly, collision probabilities in a landscape are intrinsically linked to the number of encounters an animal has with roads, and peaks in activity such as mate searching, breeding and juvenile dispersal have repeatedly been cited to explain seasonal patterns in WVC numbers (Steiner *et al*., 2014; Ryu & Kim, 2021; Raymond *et al*., 2021). Despite this, the influence of patterns of space use by animals on WVC has yet to be adequately addressed in mechanistic models.

Patterns of space use are described within the frameworks of movement ecology. Species distributions are generally divided into broad strategies comprising several patterns of space use that depend both on the biological traits of the species and the environment: sedentary populations that occupy a restricted geographical area (often referred to as home range or territory), migratory individuals who exhibit long-distance and often periodic displacements, and nomadic populations whose movements are neither confined to specific geographic areas nor predictable (Mueller & Fagan, 2008). These large-scale patterns emerge from movement at the individual level: for example, oriented movement, based on environmental cues or memory, is the basis for migratory routes and sedentary ranges (Mueller & Fagan, 2008; Van Moorter *et al*., 2009). The advances of the last decades in biologging (*e.g* animal-borne GPS collars, tags or transponders) have made it increasingly easy to acquire position data characterising these individual trajectories for a growing number of species, therefore providing a solid empirical basis to develop mathematical models of animal movement. To our knowledge however, WVC studies seldom incorporate paradigms from movement ecology, although space use is the main driver of animal encounters with roads. Therefore, we argue that modelling wildlife-vehicle collisions from a movement ecology perspective would improve our ability to understand the main factors driving WVC (Hels & Buchwald, 2001; van Langevelde & Jaarsma, 2005).

We here propose an individual-based model to simulate animal trajectories using biased and unbiased correlated random walks, as a mean of exploring the effects of road characteristics, species traits and patterns of space use on the number and probability of collisions between animals and vehicles. Our two main goals are *i)* to quantify the relative effects of several road characteristics (road sinuosity, vehicle speed and traffic volume) and species traits (animal speed, local abundance, movement patterns, foraging and dispersal behaviours) on WVC occurrences and *ii)* for a set of biologically realistic scenarios of animal movement, to identify what factors have the largest impact on wildlife-vehicle collision risks. Understanding which biological, human and environmental factors modulate the risks of wildlife-vehicle collisions is key to the selection and implementation of efficient mitigation policies such as fences, over or underpasses, and traffic signage that raises drivers’ attention (Davenport & Davenport, 2006). Our model should help to build qualitative predictions either for inter-specific studies or WVC mitigation.

## 2 Material & Methods

Following previous work on simulations for the ideal gas model (Hutchinson & Waser, 2007), we constructed a discrete-time simulation of particles movement in which all trajectories were approximated by a series of straight-line steps of constant duration. Two classes of particles were considered: vehicles and animals, each with their own speed, movement patterns and density. We conducted preliminary testing of the model by replicating the conditions under which the ideal gas model applies, and subsequently ensured that simulation results matched the model’s predictions. From there, we used biased and unbiased correlated random walk models to represent animal movement more realistically. Random walk parameter values were chosen to match real-life scenarios. The goal of the simulations was to keep track of the number of encounters between vehicles and animals per unit of time. We set a duration of 12 hours per simulation run, as a broad representation of the amount of time during which most diurnal and nocturnal terrestrial species are active (Refinetti, 1999). Two types of experiments were conducted: (1) to test the quantitative and qualitative effects of the different focal parameters (animal and vehicle speed, movement and densities) independently, we replicated the simulation for different values of the parameter under study, holding everything constant and (2) to evaluate the relative importance of each focal parameters on the number of WVC, we conducted one-at-a-time sensitivity analyses.

### 2.1 Basic simulation layout

We first assumed two separate entities, thereafter designated ‘vehicles’ and ‘animals’, moving on a 30×30km^2^ plane mapped by *x* and *y* axes. We treated the plane like a torus, meaning that an animal or a vehicle crossing the edge reappeared at the same location, but on the opposite edge of the plane (see Fig. 1). The vehicle movement followed a road defined as 0 *≤ x ≤* 30 and *y* = cos(*α*.*x*), where *α* controlled road sinuosity. We computed an index of sinuosity for each simulated road as the ratio of the road’s length to 30 (*i.e.* the x axis’ length). A sinuosity index of 1 indicated a straight road, and the index increased as the road became more and more sinuous. The number of vehicles on the road at any given time followed a Poisson distribution (Breiman, 1963) and all vehicles moved at constant speed with no intra-class variability.

**Fig. 1:**
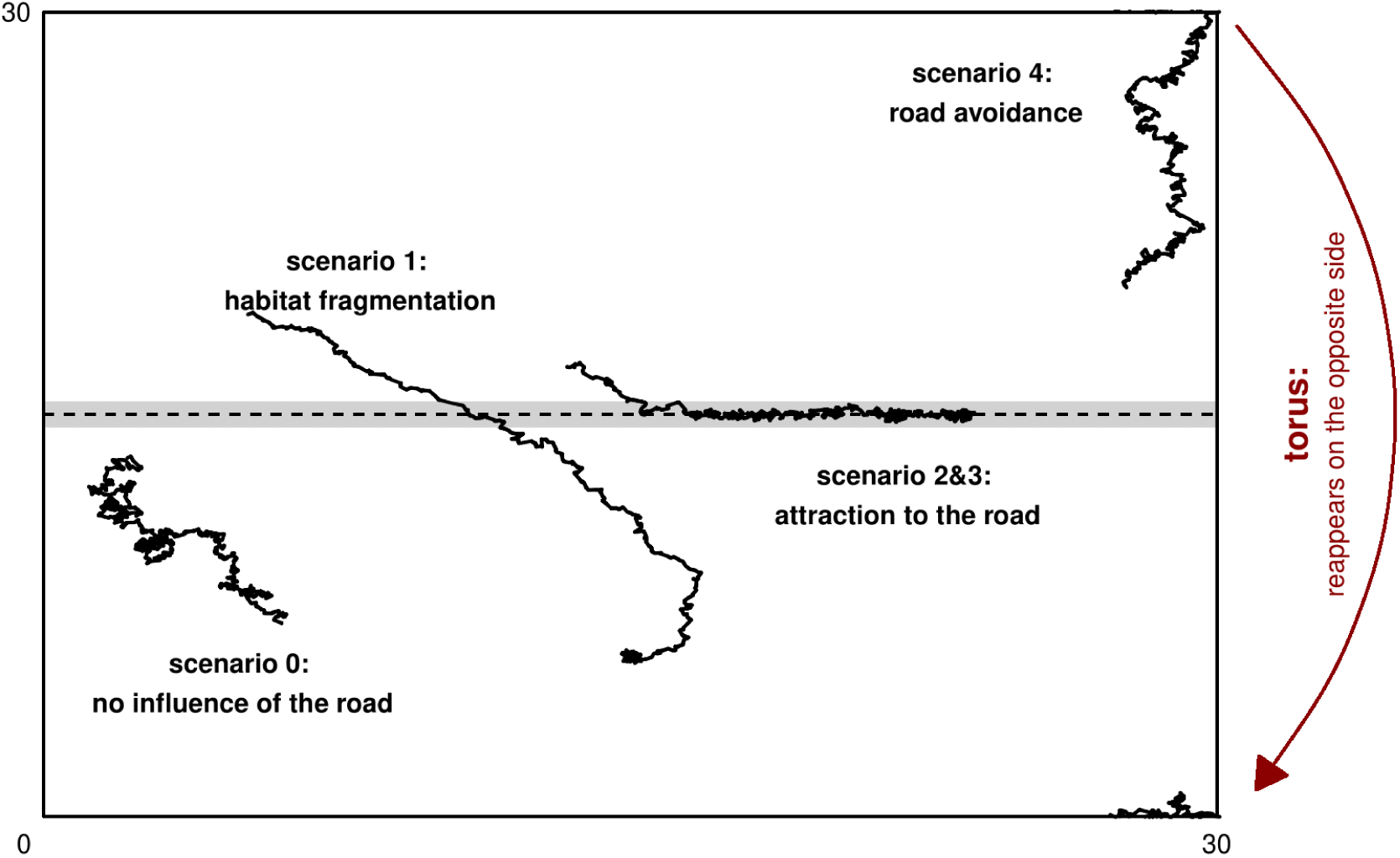
Using biased correlated random walks, 5 scenarios of animal habitat use are implemented on the 30×30km² plane to study collisions between moving animals and vehicles. Animals and vehicles that reach one of the edges of the plane will continue their walk on the opposite edge. In other words, they move on a torus and can never leave the simulation plane.

For animal movement, we used biased correlated random walks (BCRW), a combination of correlated random walks where animals have a directional persistence and therefore avoid backtracking, and biased random walks where animals are attracted to one or several centers of attraction spatially defined on the simulation plane. BCRW are highly flexible models proven to fit animal trajectories at different spatial scales for a range of terrestrial species in heterogeneous landscapes (red deer (*Cervus elaphus*): Berthelot *et al*., 2020; caribou (*Rangifer tarandus*) and bears (*Ursus arctos* and *U. maritimus*): Auger-Méthé *et al*., 2016; cactus bugs (*Chelinidea vittiger*): Schooley & Wiens, 2003).

Animals moved on the plane from time steps *t* = 0 to *T* according to a BCRW:

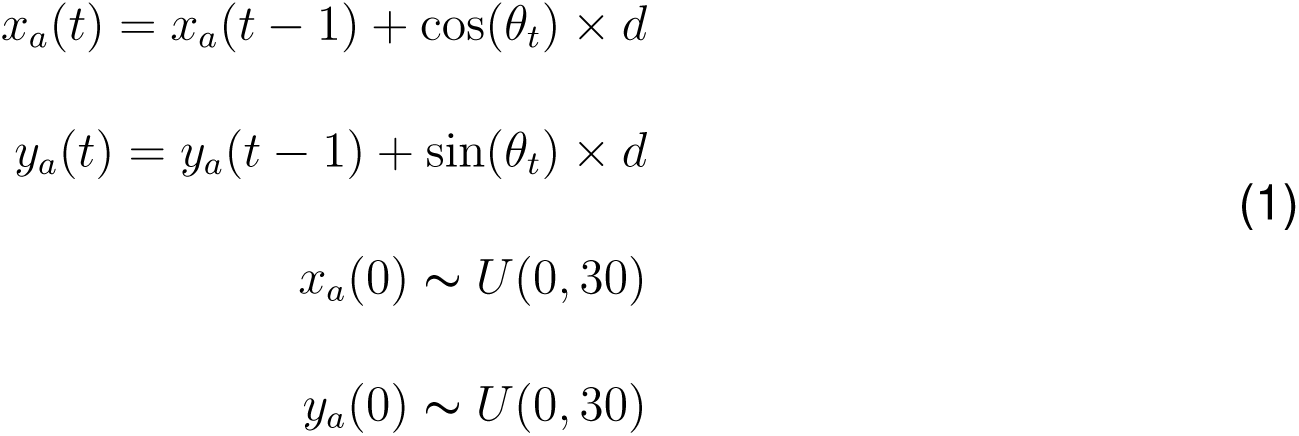

with step length *d* drawn from a zero-truncated normal distribution *N* (*µ*, .5), *µ* the mean animal speed and *U* (0, 30) the uniform distribution between 0 and 30. We computed the direction of the walk 0 *≤ θ_t_ ≤* 2*π* as the mean between the *correlated* direction 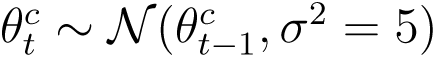, with 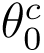 randomly drawn between 0 and 2*π*, and the *biased* direction *θ^b^*, the direction of the nearest attraction center. It resulted that the direction of the walk was the outcome of two competing processes: the *persistence*, the animal’s internal propensity to walk without backtracking in a given direction 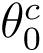, and the *attraction* toward the nearest attraction center. The relative importance of the attraction with respect to the persistence was defined by the associated weight of attraction *w* (Duchesne *et al*., 2015). Accordingly, *w* controlled the strength with which animals are attracted to the attraction center: *w* = 0 produces unbiased correlated random walks (no attraction), and *w → ∞* tends towards biased random walks. In practice, for high values of *w* the animal made a beeline towards the nearest centre of attraction because the *attraction* to the nearest center takes over the *persistence* of the correlated walk in orienting the walk. Once the animal reached the attraction center, it orbited around it, as a result of the *persistence* pulling away in its own direction while the *attraction* component brought the animal back towards the center. In consequence, high values of *w* would imply less spread-out orbiting behaviours, mimicking animals staying in close proximity of the attraction center (see Fig. S1 in supplementary material).

We defined an animal-vehicle collision as a time-step during which the distance between an animal and a vehicle was less than 5 meters (Euclidean distance). If that condition was satisfied, the animal died and stopped moving, while the car continued to move along the road. For each collision, a new animal would enter the plane in order to keep the animal density constant. We did not implement any sort of behavioural reactions from the animals to an incoming vehicle, or from the driver in the presence of an animal on the road or roadside. Given its stochastic nature, the model was run *N* = 1000 times to produce a mean collision number and estimate an individual probability of collision. The simulation used R package *Rcpp* (Eddelbuettel & Balamuta, 2018).

### 2.2 Behavioural scenarios of animal movement and space use

We considered 5 different scenarios designed to mimic real-life animal behaviours described in the literature (Fig.1).

#### 2.2.1 Scenario 0: no influence of the road

In scenario 0, the weight of attraction *w* was set at 0 resulting in animals performing (unbiased) correlated random walks (CRW). In CRW, successive steps directions are correlated, resulting in a directional bias: the animal walked forward and rarely backtracked. CRW have been used to describe the movement of several species (Mouse (*Apodemus sylvaticus*): Benhamou 1991, Reindeer (*Rangifer tarandus*): Må rell, Ball & Hofgaard 2002, Red fox (*Vulpes vulpes*), Snowshoe hare (*Lepus americanus*): Siniff & Jessen 1969). CRW imply that animals did not factor in the presence of a road or moving vehicles in their movement, which could be especially prevalent on smaller or unpaved roads (Brehme *et al*., 2013). This simple movement model also served as a null model, *i.e.* a scenario without attractive or repellent effects of road presence on animal movement.

#### 2.2.2 Scenario 1: habitat fragmentation

All animals started the simulation run at the top-left corner of the simulation plane, always above the road, while the single attraction center (*w*=1.5) was randomly located below the road. The resulting BCRW simulated a migration of individuals between patches during which they had no choice but to cross the road. This scenario could match many migrating species such as amphibians converging towards open waters for reproduction (Hels & Buchwald, 2001; Sillero, 2008), large mammals migrating to their summer or winter ranges (Avgar *et al*., 2014) and crab species moving back to sea during breeding migrations (Ryu & Kim, 2020).

#### 2.2.3 Scenarios 2 & 3: attraction to the road

Scenario 2 emerged from the frequently reported attraction to roads in many species: salt licking on roadsides by large mammalian herbivores (Fraser & Thomas, 1982), use of roads and roadsides for foraging or hunting (Diurnal raptors: Meunier *et al*., 2000; Passerines: D’Amico *et al*., 2013) or carnivores scavenging on carrions from animals hit by cars (Schwartz *et al*., 2018; Ratton *et al*., 2014). Arguably, this scenario could encompass the case of reptiles that bask on warm road surfaces (Meek, 2009). Attraction centers (*w* = 3) were placed every 300m on the road, such that animals converged towards the closest attraction center. Once an animal had reached an attraction center, it would orbit around until another attraction center in the vicinity became the new closest center, at which point the animal would leave its current attraction center to walk towards the new one. Little by little, an animal would visit several attraction centers during a simulation run, and would often walk on or in close proximity to the road while moving in-between attraction centers.

Scenario 3 is a slightly modified version of scenario 2, intended for animals with flying ability (birds, bats…) and using roads as corridors for long-range displacements, or for hunting because roads “drive” preys in the landscape (Kerth & Melber, 2009; Schwartz *et al*., 2018). Flying animals often make low-altitude flight to catch a prey or land on the road to feed (Grilo *et al*., 2014; see also the ‘leap and strike’ hunting behaviour of owls: Norberg & Norberg, 1970; Southern, 1954). To this end, we added a third dimension *z* describing the altitude of the animals to their existing (*x*,*y*) position on the plane. The altitude was a function of the distance to the nearest attraction center: *z*(*t*) = exp(0.3 *∗ ||d||*) *−* 1, *||d||* the Euclidean distance between the animal and the closest attraction center. This distribution of altitudes entailed that animals got progressively closer to the ground as they approached the attraction center. In consequence, they were only susceptible to collision when within a radius of 5m around a center of attraction.

#### 2.2.4 Scenario 4: road avoidance

The last scenario accommodated animal movement for road avoidance behaviours. Road avoidance has been reported in several species (Grizzly bear (*Ursus arctos*): Mace *et al*., 1996; Wild boar (*Sus scrofa*) and Red deer (*Cervus elaphus*): D’Amico *et al*., 2016), possibly resulting from chemical, noise pollution, or road surface avoidance (Andrews & Gibbons, 2005; Schaub *et al*., 2008; Jaeger *et al*., 2005). Building upon scenario 2, we only modified the *attraction* angle *θ^b^* to be computed as *θ^b^* + 180*^◦^*, meaning that the animal moved away from the center instead of towards it.

### 2.3 Qualitative and quantitative predictions of the model

For each of the scenarios described above, we derived qualitative and quantitative predictions for 6 parameters of interest, by running the simulation at different values of this parameter while holding everything else constant. The parameters of interest were: animal mean speed, vehicle speed, animal density, traffic volume, weight of attraction centers *w* (scenarios 1 through 4 only) and road sinuosity (Table 2).

**Table 2:** The six parameters of interest, relating to animal and vehicle movement, and that appear to be pivotal factors driving collision numbers according to most studies (Table 1). While the effects of one parameter on collisions are being explored, all other parameters assume their baseline values. To produce quantitative and qualitative predictions, we explored for each parameter a range of values contained within biologically relevant intervals.

|  | Parameter of interest | Baseline value | Explored range |
| --- | --- | --- | --- |
| ANIMAL | Mean speed (km.h<sup>-1</sup>) | 5 | 0.1 - 30 |
|  | Density (ind.km<sup>-2</sup>) | 0.016 | 0.001 - 0.03 |
|  | *w* | 5 (*scenario 1*); 3 (*scenarios 2-4*) | 1 - 20 |
| VEHICLE | Speed (km.h<sup>-1</sup>) | 80 | 30 - 300 |
|  | Traffic volume (vehicle.h<sup>-1</sup>) | 30 | 10 - 250 |
|  | Road sinuosity | 1 (*straight road*) | 1 - 3.5 |

Mean animal travel speed values covered the observed speed range of most terrestrial species (from slowest to fastest, migrating amphibians: Hels & Buchwald 2001, foraging badgers (*Meles meles*): San *et al*. 2007, migrating ungulates: Berger *et al*. 2006; Singh & Ericsson 2014). Considering that this range is wide, we extracted mean travel speed estimations for several species from the literature (Rowcliffe *et al*., 2016; Farley *et al*., 1993) and set our baseline at the mean value of 5 km.h*^−^*^1^. Vehicle speed encompassed slow vehicles up to high-speed trains, and the baseline was set at the current speed limit on main roads in France (80 km.h*^−^*^1^). Local animal densities, traffic volume ranges and baselines were mostly dictated by computational limitations, although the baseline of 250 vehicles.hour*^−^*^1^ is representative of average sized roads in France (French Ministry of Ecological Transition, 2022).

The weight of attraction *w* in BCRW scenarios was a dimensionless unit ranging from 1 (attraction to centers is equal to the *correlated* direction of the walk, spread-out orbiting behaviour around attraction center) to 20 (when attraction was 20 times more important in walk orientation, animals walked in nearly straight paths toward the attraction center and stayed mostly in place once it was reached). Baseline values for *w* were scenario-dependent as *w* = 5 for scenario 1 ensured that over 90% of animals cross the road in less than 12 hours and *w* = 3 for scenarios 2 and 3 ensured that animals would effectively visit several attraction centers during the simulation run. Finally, road sinuosity was investigated following observed sinuosities which are publicly available for the road network of Ireland (Transport Infrastructure Ireland, 2022).

Two types of results were extracted from simulation runs: the mean number of collisions recorded over the course of a 12 hours run, and the individual probability of collision for animals. Considering that new individuals replaced dead animals, not all animals were present from the start of the simulation run and collision risk was not constant across all animals. Consequently, we estimated the individual probability of collision from a logistic regression (0: alive, 1: hit by a vehicle) as a function of the parameter under study, and where the time of arrival in the simulation was entered as an offset predictive variable (Agresti & Coull, 2002).

### 2.4 Sensitivity analyses of collision number

Sensitivity analyses were conducted on each scenario, following a *one-at-a-time* approach, meaning that we measured the variation in collision numbers while keeping all input factors fixed except the one that is being perturbed (Saltelli *et al*., 2019). In practice, we computed the mean number of collisions *y* for factor of interest *x* at a reference value *x*_1_, and after 10% increase in value (denoted *x*_2_). To explore possible non-linearity, we repeated this for several reference values *x*_1_ within the relevant range of *x* (see section 2.3 for parameter ranges). We then defined the sensitivity measure as *OR_x_*_1_ = *y_x_*_2_ */y_x_*_1_, *i.e.* the odd ratio of the collision probabilities that results from a 10% increase of parameter *x* at reference value *x*_1_. For example, an *OR_x_*_1_ of 1 indicated no change in the odds of collisions between parameter values *x*_1_ and *x*_1_+10%, while *OR_x_*_1_ *>* 1 indicated greater odds of collision after a 10% increase in *x*_1_. The overall strength of the influence of parameter *x* on the collision number in its relevant range was defined by how far the mean of the ratios (OR*_mean_*) for all tested values of *x* deviated from 1 (*e.g.* OR*_mean_* = 1 indicated no overall influence of factor *x* on collision numbers).

## 3 Results

### 3.1 Qualitative and quantitative model predictions

Applying baseline values to all parameters of interest, mean values ranged from 0.16 to 17.2 animal-vehicle collisions over 12 hours (individual probabilities of collision for animals: 0.4% - 45.6%). As expected, the movement of animals had major effects on the probability for an animal to get hit by a car (Fig.2). With baseline parameter values, animals attracted to the road (scenarios 2 and 3) were hit more often than in any other scenario, with an increase in odds of collision for terrestrial animals (scenario 2) compared to flying animals such as birds and bats (scenario 3). Compared to the null model (scenario 0, roaming with no attraction/avoidance of the road), migrating animals crossing the road exactly once (scenario 1) were hit more often, while the odds of collisions for active road avoidance (scenario 4) were reduced. This ranking between scenarios was not always maintained for other values of the tested parameters.

**Fig. 2:**
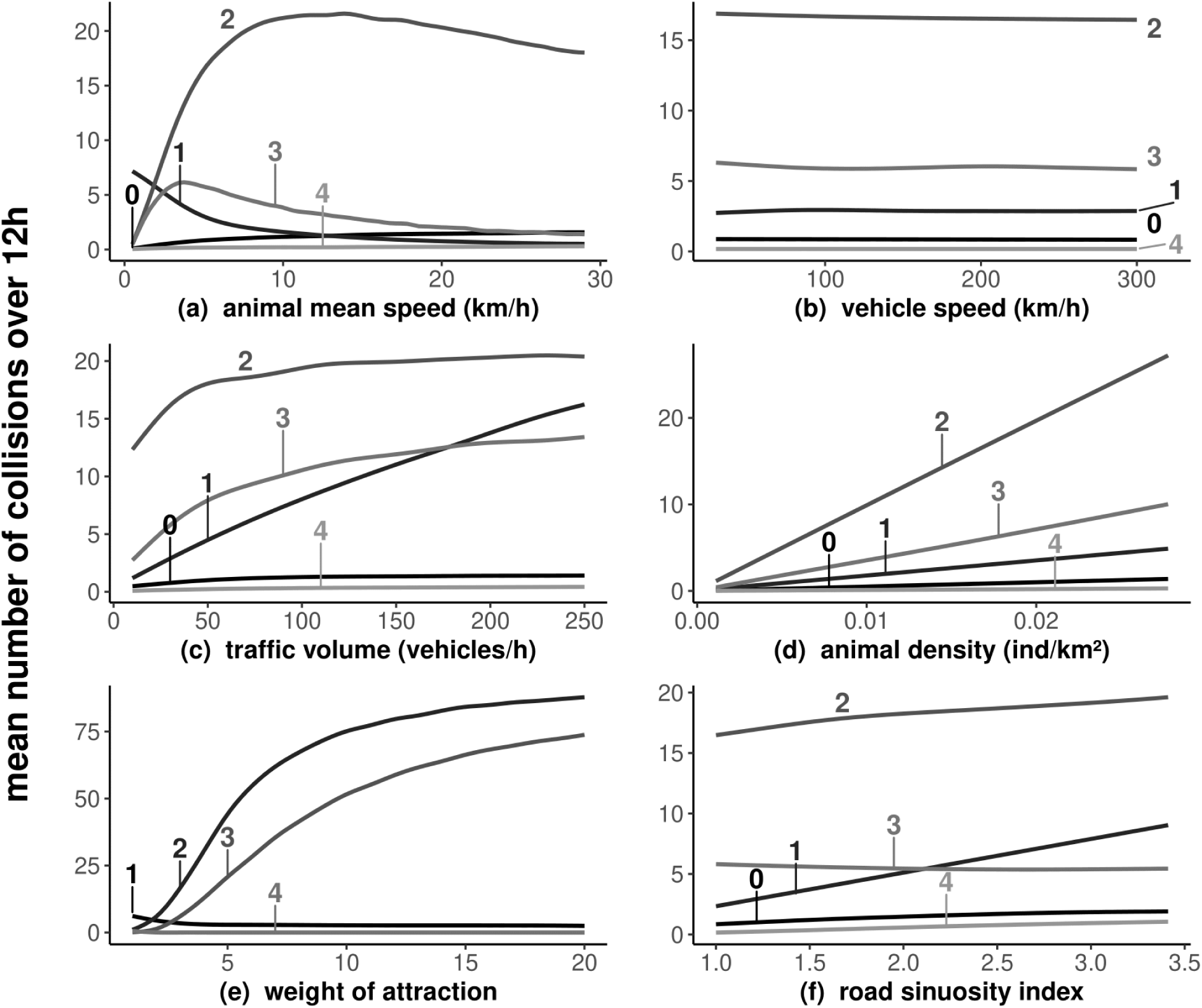
Mean number of collisions. For each scenario, the total number of collisions occurring within a 12-hours simulation run (mean number over 1000 runs). The weight of attraction *w* has no meaning for (non biased) correlated random walks and is therefore not considered for scenario 0.

Overall, parameters response functions had similar shapes whether the outcome was measured as the mean number of collisions or as the individual probability of collision, with the exception of animal density. Vehicle speed had no effect on collision numbers and probabilities in any of the scenarios (Fig.2b and 3b). Animal density was positively and linearly associated with collision numbers, which translated into a constant individual probability of collision no matter the local abundance in animals (Fig.2d and 3d). The influence of the remaining parameters (animal speed, weight of attraction, traffic volume and road sinuosity) was non-linear and/or scenario-dependent.

**Fig. 3:**
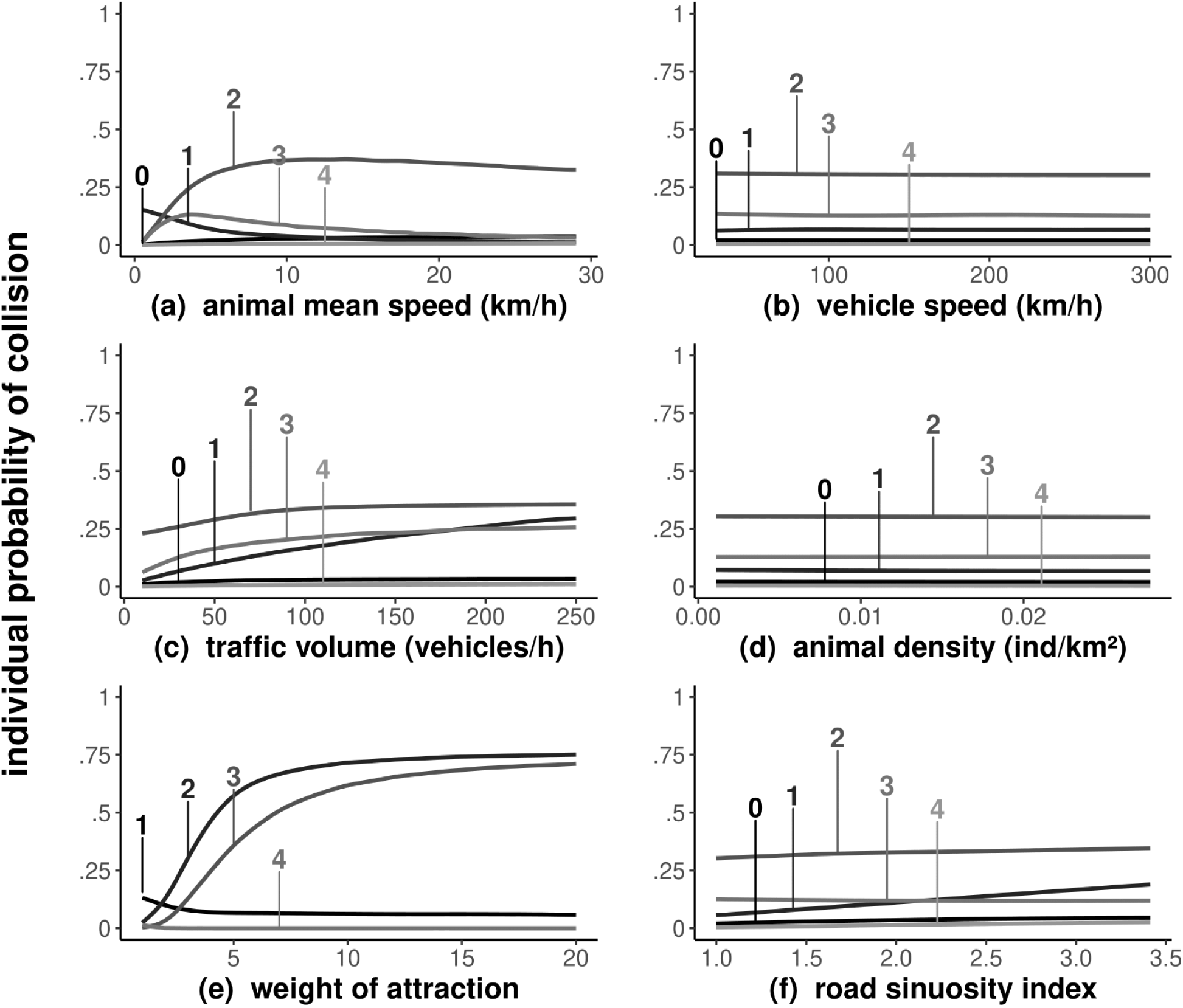
Individual probability of collision. For each scenario, the individual probability of collision estimated from 6 separate logistic regressions in which the studied parameter was used as a response variable and the individual time of exposure to vehicles was used as an offset.

Traveling faster reduced collision numbers and probabilities for migrating animals (scenario 1) but had the opposite effect on roaming and road-avoiding animals (scenarios 0 and 4). In scenarios with road-attracted animals, collision occurrences increased with mean animal speed up to moderate speed values, after which the relation was reversed (Fig.2a, 3a). Higher volumes of traffic on the road increased collision numbers and probabilities although scenarios 0, 2, 3 and 4 present inflexion points after which the increase in collision occurrences slowed down, which was not observed within the explored range of traffic volume for scenario 1 (Fig.2c, 3c). Similarly, increasingly sinuous roads had moderate to low positive effects on collision number and probabilities in all but one scenarios: scenario 3 yielded no apparent effect of road sinuosity (Fig.2f, 3f). Finally, increasing weights of attraction (or *repulsion* for scenario 4) led to more collisions (and higher individual probabilities) in scenarios 2 and 3 with inflexion points around *w*=5, but increasing *w* values were associated to less collisions and lower probabilities of WVC in scenarios 1 and 4 (Fig.2e, 3e).

### 3.2 Sensitivity analyses

For roaming animals (scenario 0), the most influential parameter was the animal density (OR*_mean_*=1.091) followed by animal speed (OR*_mean_*=1.032). In all other scenarios (scenarios 1, 2, 3 and 4), the weight of attraction *w* had the largest influenced (respectively OR*_mean_*=0.888, 2.019, 1.155 and 0.843) followed by animal density (respect. OR*_mean_*= 1.098, 1.095, 1.101 and 1.108). The least influential parameters was traffic volume for scenario 0 (OR*_mean_*=0.996), vehicle speed for scenarios 1 and 3 (respect. OR*_mean_*=1.01, 1.006) and road sinuosity for scenarios 2 and 4 (respect. OR*_mean_*=0.998, 0.993). Using OR*_mean_* is, however, a simplified view of the non-linear effect of parameters, as it was sometimes pulled in one direction by one or a few extreme OR values. For example, some parameters such as the weight of attraction/repulsion generally had a substantial influence on collision numbers at lower values (*w* = 1) while higher values had OR values closer to 1 (Fig.4).

**Fig. 4:**
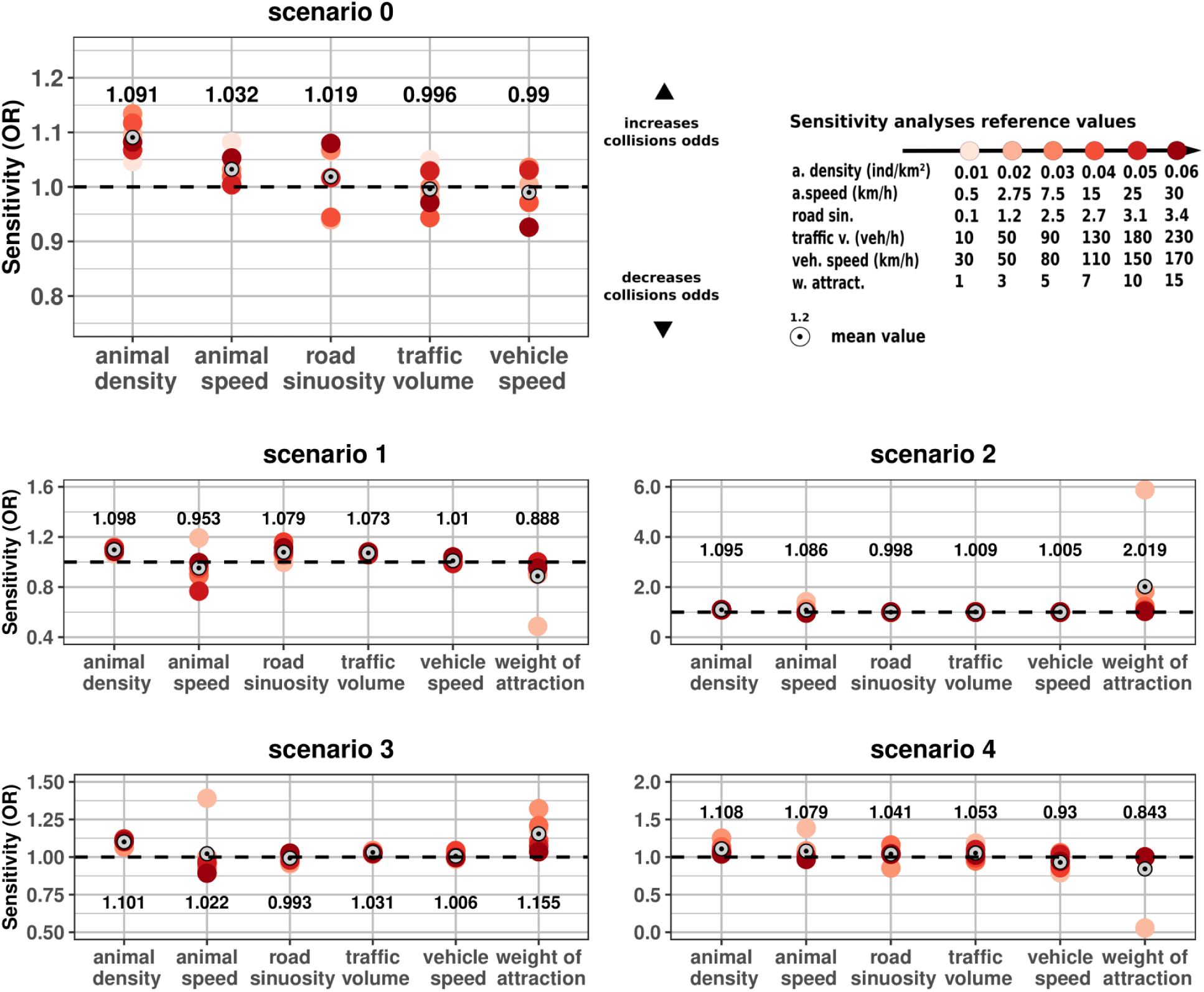
Sensitivity analyses. We compute the odds ratios (OR) of the mean number of collisions for a 10% increase in a given parameter reference value. For example, the darkest circle (*•*) under “traffic volume” is the ratio of the mean number of collisions with 230 + 10% = 253 vehicles/hour over the mean number of collisions with 230 vehicles/hour. An OR *>* 1 shows that an increase in traffic volume has increased the number of collisions (and conversely for OR *<* 1). The further from 1 the mean is, the more influential the parameter is on collision numbers.

## 4 Discussion

Road mortality of wildlife is a pervasive and widespread cause of biodiversity loss across the world that should be seriously considered in wildlife management and conservation programs. In spite of its ecological importance and relevance, we are still lacking a strong formalisation of WVC to grasp what are its main biological, ecological and anthropogenic determinants. By combining earlier encounter models with the current knowledge about movement ecology, our work should help at formulating qualitative predictions for reducing the number of WVC or to predict what species should be at greater risks of being hit by a vehicle.

Our simulations show how collision numbers and probabilities differ markedly with the movement behaviour of animals, such as migration or ranging, or with its locomotion (terrestrial *vs.* flight). We suggest this contrast in mortality risk between individuals with different movement types could lead to selection forces on animal habitat use with regards to road avoidance or attraction. In practice, the knowledge about animal movement of species should be an essential part of WVC studies as it seems the most influential variable on WVC occurrence compared to any changes in vehicle speed or road sinuosity for instance.

### 4.1 Effects of animal movement on WVC risks

WVC broadly results from two competing mechanisms: the number of times an individual will cross or stay close to a road, and the time spent at risk of collision when on the road while crossing, foraging or basking. Both mechanisms are modulated by patterns of space use (in our simulation, the different scenarios of behaviour), and their modulations through the weight of attraction/repulsion *w* and the speed at which animals travel. Obviously, individuals actively avoiding roads (scenario 4) should be the least at risk of collision with a vehicle and, conversely, terrestrial individuals attracted to the road for foraging or basking (scenario 2) should be the most at risk. This is in line, for example, with Western green lizards (*Lacerta bilineata*) being roadkilled in higher proportions than the other sympatric lizard species that do not bask on roads (Meek, 2009). Flying carrion feeders (mostly diurnal and nocturnal raptors) are less likely to be killed than their terrestrial counterparts, a difference that may be further widened by some species’ cognition and ability to successfully evade oncoming vehicles, as is the case with corvids (Mukherjee *et al*., 2013; Lima *et al*., 2015). Nevertheless, we show here that other parameters of both animal and vehicle movement have the ability to modulate these conclusions. For example, slow moving animals (less than 15km.h*^−^*^1^) were more often hit when migrating (scenario 1) than roaming freely (scenario 0), but the reverse was true for faster animals (Fig.2a).

By mimicking the presence of attractive or repulsive areas for animals in the landscape, the attraction/repulsion weight (*w*) ultimately controls the time spent in the vicinity of the road, which is positively correlated to collision numbers and probabilities. The relationship between *w* and the time spent on the road is easily interpreted for scenarios of road attraction (scenarios 2 and 3, *w* is positively correlated to time on road surface) and avoidance (scenario 4, *w* value is negatively correlated to time on road surface). When animals travel from a departure point to an attraction center (scenario 1), the time spent crossing is negatively correlated to *w*: high *w* values means straight trajectories between departure and attraction center, and thus less time on the road.

Collision risks in animals roaming freely in a landscape (scenario 0, no *w* variable) are dictated by their speed: the faster the animal, the more it explores its environment during a simulation run and thus encounter roads more often, even though each crossing becomes concurrently less dangerous as fast animals cross the road in less time (see Fig.S2 in supplementary material). For the same reason, when animals cross the road only once (scenario 1), we would also expect the animal travel speed to be a key variable in decreasing collision numbers. However, this was not reflected by *OR_mean_* in the sensitivity analyses, the most likely explanation being that the mean sensitivity *OR_mean_* for travel speed in scenario 1 is pulled up by the odds ratio for very low travel speeds (odds ratio *OR*_0.5_ comparing collisions at 0.5 and 0.55 km.h*^−^*^1^) where more animals would cover the distance between their departure point and the road during the simulation when traveling at 0.55 km.h*^−^*^1^ (Fig.4). Nevertheless, these results demonstrate that the cumulative time spent on or near roads by animals, through repeated road encounters and/or a long time spent on the road at each encounter (modulated here by patterns of space use, animal speed and biased random walk parameter *w*), is the most influential parameter driving collisions probabilities for an animal, and has more importance than road characteristics such as traffic volume or speed limits.

These findings are consistent with studies exploring the relationship between landscape features and roadkill hot-spots: factors such as the presence of attractive habitats on roadsides as well as a high connectivity landscapes conducive to wide ranges of movement are often linked to high road mortality (Table 1, see also de Freitas *et al*. 2013; de Freitas *et al*. 2015). Similarly, the temporal distribution of roadkill is dictated by the level of activity, and periods of high mobility such as rut, mating or migrating seasons are especially risky for animals (Table 1). Larger home ranges and spread-out exploratory behaviours contribute to higher fitness for individuals by increasing foraging opportunities (Andersson, 1978), but the presence of roads (also applicable to aircrafts, ships or trains) in a landscape lead to costs in the form of road mortality and selection pressure may lead to road avoidance (Meisingset *et al*., 2013; Jaeger *et al*., 2005; Husby & Husby, 2014).

### 4.2 Effect of animal density on WVC risks

The impact of spatial and temporal variations in animals density has been repeatedly reported as a significant driver of WVC (Joyce & Mahoney, 2001; Taylor *et al*., 2010; Saint-Andrieux *et al*., 2020). Some authors suggested that the number of carcasses on roads could serve as an index of animal abundance (Seiler & Helldin, 2006; George *et al*., 2011). We show that this relationship is linear but strongly modulated by animal movement (Fig.2): compared to animals with no attraction or avoidance towards roads, the slope of the relation between local density and roadkill number is 8 times greater for terrestrial animals attracted to roads and about 70 times smaller when there is road avoidance. When dealing different species, or with large temporal or spatial scales in which animal movement might be modified by biological seasons and/or landscape features, WVC counts are meaningless, unless the average movement of animals is precisely known. Furthermore, all slopes deviates from isometry (*β <* 1), suggesting that the number of casualties on the roads underestimates high population abundances.

Animal density has an impact on collision numbers, but not on the probability of collision for the individual: WVC are coincidental and have no upper threshold for the number of roadkill, contrary to, for example, a predator looking to feed on a set number of prey. Because of this, there is no mechanism of dilution, or “safety in numbers”, in which we might expect a lower collision risk for the individual in a large group of conspecifics (Lehtonen & Jaatinen, 2016). We need to mention that changes in local density have documented effects on individual behaviour which were not explored in the model, and that will in turn affect movement (Kjellander *et al*., 2004; Dahle & Swenson, 2003; Sword, 2005).

### 4.3 Effects of traffic volume on WVC risks

A general consensus arises in the literature on the number of WVCs increases with road traffic (Fahrig *et al*., 1995; Rosen & Lowe, 1994; Inbar & Mayer, 1999; Joyce & Mahoney, 2001). This correlation is, however, much discussed with some authors proposing a linear relationship (Fahrig *et al*., 1995); and others a *barrier effect* (high-traffic roads having fewer WVCs than less frequented roads because animals avoid the noise and other disturbances they generate: Seiler & Helldin, 2006; Clevenger *et al*., 2002; van Langevelde *et al*., 2009). Others explained the lower-than-expected number of collisions reported on some high-traffic roads by a depletion of local animal populations due to roadkill (Ascensão *et al*., 2019). Those arguments may be unnecessary given our results: the theoretical relationship between traffic and WVCs can be non-linear even in the absence of traffic-dependent road avoidance or source-sink dynamics, and roads with heavy traffic would not necessarily have higher collision rates than intermediate roads.

Whatever assumption we make on animal movement, the number and the individual probability of WVC grow asymptotically within the explored range of traffic (Fig. 2 and 3). This observation agrees with the mechanistic model of WVCs by Hels & Buchwald (2001) that too predicts a logarithmic relationship between traffic volume and WVC occurrence. A noticeable difference between the two models is that while the probability of dying animals asymptotically approaches 1 for Hels & Buchwald’s model, the asymptotic number and probability of WVC lie below a 100% in all our simulations. In one case no asymptote is visible (scenario 1), which possibly results from the limited range of traffic volume we considered due to computational limitations (major highways worldwide can see up to 15 000 vehicles per hour).

### 4.4 Effects of vehicle speed on WVC risks

In our model, the speed of the moving vehicles is unrelated to the number and individual risk of WVC over the simulated range of speed (0–300 km.h*^−^*^1^, see Fig. 2 and 3). At first sight, a lack of effect of vehicle speed may appear at odds with the previous theoretical models suitable for WVC but it is not: in the ideal gas model (Hutchinson & Waser, 2007), the equations for encounters feature the *length of the path* of the molecule (*speed × time spent moving*) rather than the speed itself. In our model, the distance travelled by a vehicle is constant: they move from one end of the road to the other, and a higher speed does not equate with a longer distance, therefore speed does not increase WVC risks. In contrast, on the subject of the animal’s speed, when higher speed means longer travel distances (for scenarios 0 and 4; and scenarios 1, 2 and 3 are more complex as the movement is also directed), it leads to higher WVC occurrences and risks. In addition, the travelled distance by vehicles was increased by simulating more tortuous roads (high sinuosity index), and the number and probabilities of collision for animals increased as well (Fig.2), as expected under the ideal gas framework.

For flying carrion feeders (scenario 3) only, sinuosity is not or slightly negatively correlated to WVC numbers and individual probabilities. We do not have a satisfactory answer to this, but suggest that the euclidean distance between two attraction centers placed on the road changes with road sinuosity and could impact (with our method of simulating flight) the altitude and therefore the collision risk, which is why terrestrial animals (scenario 2) are not impacted. Road sinuosity can also be treated as a measure of the density of roads, *i.e.* the surface of road per unit of area, where sinuous roads equate to a higher density than straight roads. In that sense, we show that the density of roads is positively correlated to WVC number and risks.

Both vehicle speed and road sinuosity are among the least influential factors across all space use patterns: we would expect that fast-moving vehicles (*e.g.* high speed trains)to be no more dangerous to wildlife than slow vehicles. Empirically, speed is regarded as a significant component of WVCs: the faster the vehicles, the more likely is a collision with an animal (Table 1, see also Farmer & Brooks 2012; Hobday & Minstrell 2008). This discrepancy can be explained: fast-moving vehicles equates to reduced reaction time for both animals and drivers (Lima *et al*., 2015) and vehicles moving at high speeds are most of the time traveling on multi-lane highways with heavy traffic. The reported association between WVCs and vehicle speed may therefore be confounded with unsuccessful collision avoidance, traffic density (that we find to be a strong driver of WVC) or road width (in relation to the time spent crossing the road).

### 4.5 Model limits

We find that the time spent on the road is the critical factor driving WVC number and probabilities independently of space use patterns, and consequently behaviours we did not implement but that modify the time the animal spends on the road surface (road avoidance, pausing on the road, speeding away from vehicles, evading oncoming vehicles, see Jacobson *et al*. 2016; Lima *et al*. 2015) are likely to be key components of WVC occurrences. For example, the implementation of a *barrier effect* (avoidance of roads proportional to road traffic volume) would have predictable effects on WVC occurrences: we know that the strength of avoidance is negatively correlated to the number and probabilities of WVC (Fig.2 and Fig.3, scenario 4), which indicates that traffic-dependent road avoidance in animals roaming, migrating or foraging on the road (respectively scenarios 0, 1, 2 and 3) will decrease the number and probabilities of collisions proportionally to the volume of traffic. This would lead to the relation between traffic volume and WVC numbers to be an inverted U-shaped curve where roads of intermediate traffic volume are more dangerous than high-traffic and low-traffic roads, as conceptualized by Seiler (2006) and Jacobson *et al*. (2016). We can predict selection pressure on these behaviours will be strong for populations living in the proximity of roads, as documented in the example of road-adjacent cliff swallow populations being selected for longer wings, allowing for better chances of escaping oncoming vehicles (Brown & Bomberger Brown, 2013).

The main focus of this work was the movement of the animals, and we reduced “collisions” to the number of times an animal and a vehicle were located less than 5 meters away from each other. This will likely overestimate collision probabilities, especially for smaller-bodied species that can easily fit in-between car tyres (Hels & Buchwald, 2001; van Langevelde & Jaarsma, 2005). This is a classical simplification from the ideal gas model that we expect will not alter the general patterns we report but simply move all the curves down. Finally, we designed the simulation to have constant animal density, which does not allow for population depletion dynamics (justified by the limited temporal scale of 12 hours per simulation run). Based on our conclusions, the total number of roadkill would decrease linearly with population decline but have no impact on individual probabilities of collision (Fig.2 and Fig.3).

### 4.6 Practical implications

#### 4.6.1 Investigating roadkill patterns using movement data

The theoretical framework developed here can direct future research on roadkill spatial and temporal patterns. We have shown that collision risks in a landscape for species are highly dependant on their movement characteristics: for example, WVC in species with a home range that are not attracted nor avoiding roads (described by scenario 0) should be best predicted by the speed of movement of the individuals (Fig.4, scenario 0). Faster travel speeds for the individuals should lead to increased collision risks with vehicles (Fig.2a). In section 3 of the supplementary material, we use the case of the European roe deer *Capreolus Capreolus*, whose movements fit the scenario 0 of our model (Saïd *et al*., 2009). Roe deer movement is season-dependant, with males having distinct peaks in activity during the mating season (March to August, Krop-Benesch *et al*. 2013). In figure S3a-b, we show the seasonal increases in male roe deer speed in adults and sub-adults from the GPS data of individuals tracked during in the bavarian forest, Germany. Temporal collision patterns for this species are also known to peak during spring (Steiner *et al*., 2021; Mayer *et al*., 2021). We show roadkill reports from the citizen-science database (Faune-Auvergne-Rhône-Alpes) over a period of 12 years (2010 to 2022) in figure S3c. As predicted, seasonal increases in roe deer speed correlate to increases in roadkill reports. Efforts should be made at connecting more often movement ecology to roadkill research to understand the inter-specific and spatio-temporal variations in WVC.

#### 4.6.2 Roadkill mitigation

In this study, we demonstrate that animal movement characteristics are far more important than road characteristics in managing roadkill. More precisely, the time spent on the road by the animal is the main contributing factor to WVC occurrences, and total roadkill numbers depend on animal density rather than traffic volume or speed. In practice, mitigation measure should therefore focus on keeping animals away from the road by prioritizing fencing, or over-passes and other crossing structures for wildlife over driver-targeted measures such as signage and reduced traffic flow. Crossing structures can also restore connectivity in landscapes where roads represent a significant barrier to movement (Grilo *et al*., 2011). For species that forage on roads surfaces, we suggest that reducing the attractiveness of the road by removing carrion, salt pools and limiting the presence of preys (such as rodents) on roadsides is the most efficient solution. For example, decommissioning roadside salt pools has been shown to reduce the time moose spend on the road and could in theory reduce moose-vehicle collisions by up to 49% (Rea *et al*., 2021; Grosman *et al*., 2009). Mitigation policies should also be species-specific (Teixeira *et al*., 2013; Saint-Andrieux *et al*., 2020). Characteristics such as large large home ranges, wide diet and habitat breadths will belong to species with a particularly high-risk of WVC, as the animals that explored more of the simulation plane (scenario 0, high animal speed) were more often killed than those that stayed in smaller areas (González-Suárez *et al*., 2018; Grilo *et al*., 2020; Ford & Fahrig, 2007).

### 4.7 Conclusion

Empirical studies focusing on animal behaviour and movement in addition to other biological traits, road-side habitat type and road characteristics can give valuable insights as to why some species are more often killed on roads than others. We show here that animal behaviour in the presence of roads is a key component of roadkill occurrences, and that understanding space use patterns can give insights on the vulnerability of a species to roadkill. Although this model is not intended to provide quantitative predictions of roadkill for particular species, we show that future predictive models should incorporate components of movement ecology to properly address road mortality. Animal behaviours in the proximity of transport infrastructures could also be of interest in evolutionary studies, an aspect of road ecology that is lacking so far in the literature (Brady & Richardson, 2017).

## Supporting information

Model details

## Acknowledgments

This work was performed using the computing facilities of the CC LBBE/PRABI. We thank Simon Benhamou, Jean-Michel Gaillard and Marion Valeix for comments on previous drafts of this work.

## Author Contributions

Annaë lle Bénard: coding work and analysis of the data generated, lead writing of the manuscript, equal contribution to the methodology. Thierry Lengagne and Christophe Bonenfant: equal contribution to the methodology, writing, reviewing and editing of the manuscript. All authors contributed critically to the drafts and gave final approval for publication.

### Conflict of Interest

All authors declare that they have no conflicts of interest.

## Notes

### Competing Interest Statement

The authors have declared no competing interest.

### Summary of Updates

Add empirical results on roe deer

