## Supplementary material for "A biologically realistic model to predict wildlife-vehicle collision risks": Model details

### 1 | Supplementary Material

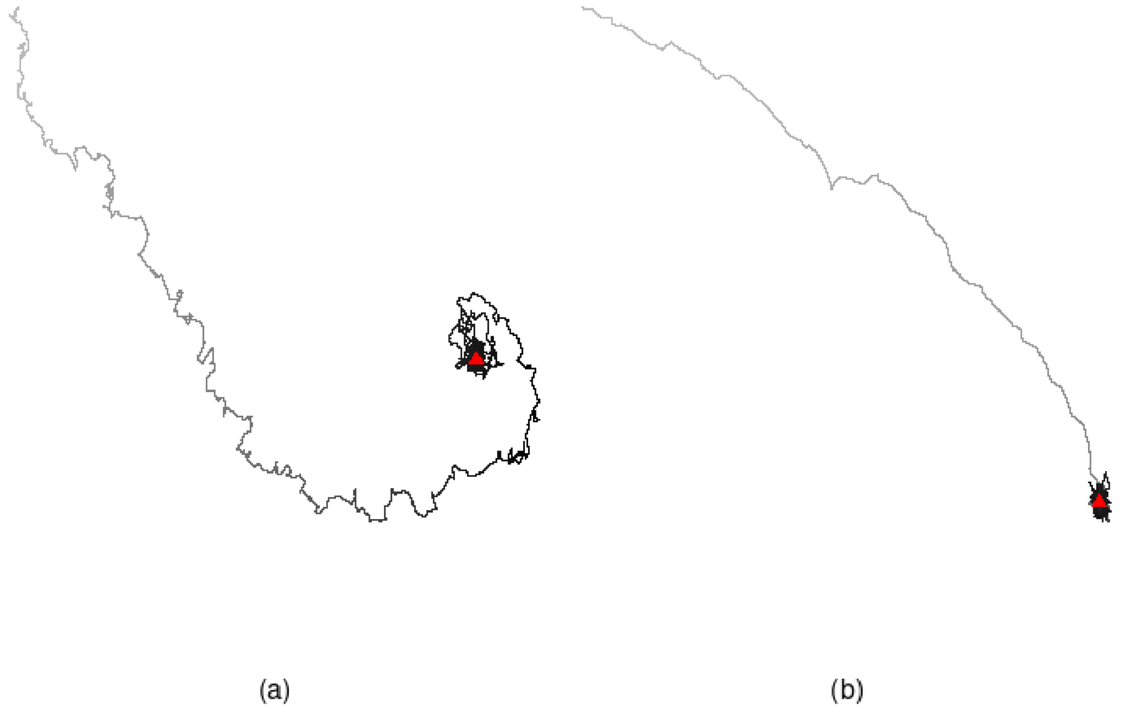


Fig. S1: The animals' movements are simulated using biased correlated random walks. The orientation of the walks is computed for each time step as a weighted mean between the persistence (correlated random walk) and the directional bias towards a center of attraction (▲). (a): the weight of attraction is $w$=3. (b): the weight of the attraction is $w$=20. The higher the weight $w$ of the directional bias, the quicker the animal reaches the center. Once the attraction center is reached, the animal will orbit around it indefinitely.

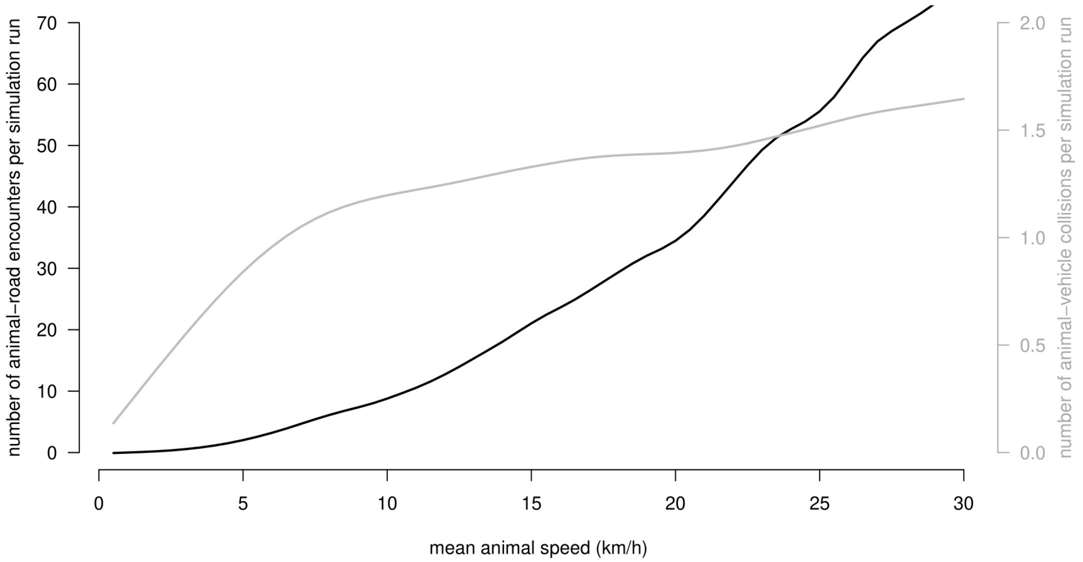


Fig. S2: For animals roaming freely (scenario 0), we retrieve from the wildlife-vehicle collisions simulation both the mean number of animal-road encounters (black line) and the number of actual animal-vehicle collisions (grey line) per simulation run. The distance each animal travels during each run increases with animal speed, leading to more animals encountering the road more often. Simultaneously, faster animals decrease their probability of collision per road crossing as they cross the road in less time. As a result, the number of collisions does not increase with the same rate as the number of road encounters: increasing the mean animal speed produces more but less dangerous road encounters for animals.

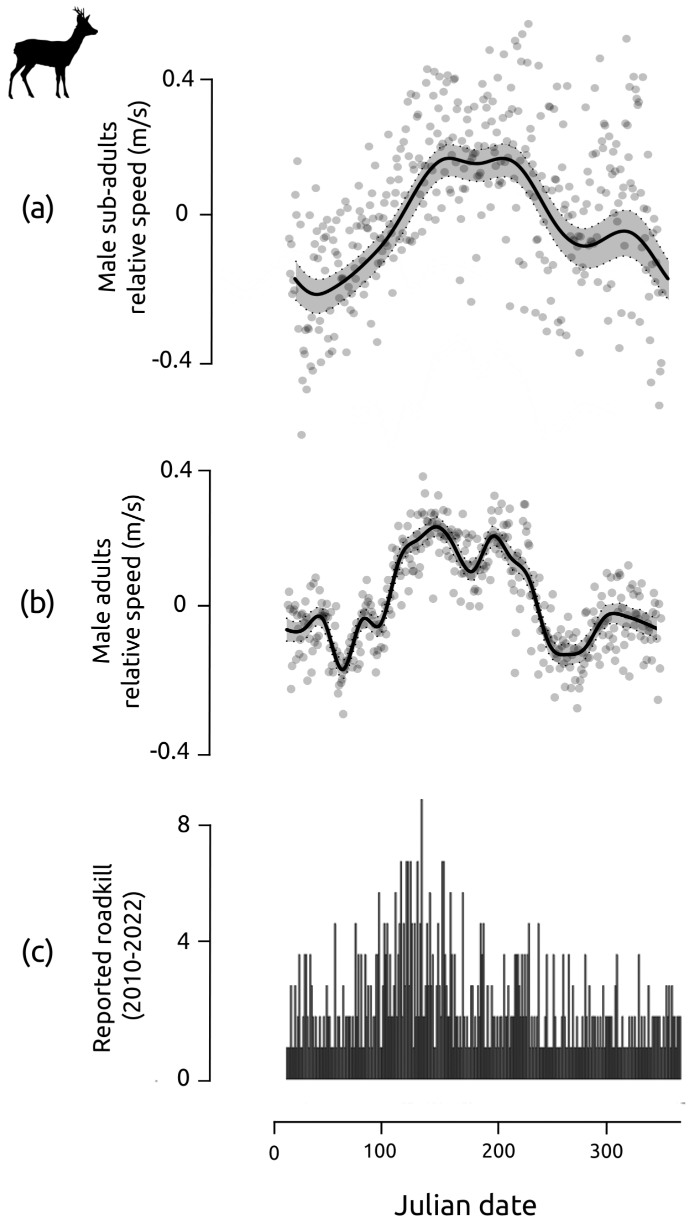


Fig. S3: Seasonal patterns in male individuals' speed (deviation to the mean, a,b) and reported roadkill (c) for the European roe deer *Capreolus capreolus*. We obtained 672 roadkill data from the citizen science database Faune-Auvergne-Rhône-Alpes (fauneauvergnerhonealpes.org), between 2010 and 2022 in the Auvergne-Rhône-Alpes region of France. The speed for male roe deer (adults and sub-adults) was measured on GPS-tracked individuals during in the Bavarian forest (Germany). Male roe deer have distinct peaks in speed and activity during the mating season (March to August). Increases in individual speed correlate to increases in the number of roe deer-vehicles collisions reported, as predicted by the wildlife-vehicle model (Fig. 2&4, scenario 0).
